# Behavioral repeatability and foraging performance in wild free-flying nectarivorous bats (*Glossophaga commissarisi*)

**DOI:** 10.1101/309971

**Authors:** Vladislav Nachev, York Winter

## Abstract

Animal individuals show patterns of behavior that are stable within individuals but different among individuals. Such individual differences are potentially associated with differences in foraging efficiency and in fitness. Furthermore, behavioral responses may be correlated in specific suites of so called behavioral syndromes that are consistent across different contexts and with time. Here we present a field investigation on individual differences between wild, free-flying nectarivorous bats (*Glossophaga commissarisi*) in the foraging context. We further investigated how individual differences effect foraging performance, and we examined their interdependence within hypothesized behavioral syndrome structures. Free-ranging bats were individually identified as they visited an array of 24 artificial flowers with nectar of high or low sugar concentration. We found that three behavioral measures of foraging behavior were individually stable over the two-month observation period. We investigated the link between individual behavioral measures and measures of foraging performance using generalized linear mixed models. Individual measures of foraging performance showed significant repeatability, and we found evidence that bats making more visits per bout tend to be slower in learning to avoid unprofitable flowers. We used a multi-response generalized linear mixed model to estimate between-individual correlations and compare hypothesized syndrome structures. There were no clear patterns of between-individual correlations among the behavioral measures in our study, despite the measures exhibiting significant repeatability. This may indicate that foraging performance depends on multiple individual behavior dimensions that are not adequately described by simple models of behavior syndromes.

## INTRODUCTION

Findings from the diverse fields of comparative behavioral biology, neurobiology, and psychology (Gosling 2001; Sih, Bell, and Johnson 2004; Dingemanse and Réale 2005; Bell 2007; Réale et al. 2007; Roche, Careau, and Binning 2016) demonstrate that intra-specific behavioral variation is sometimes maintained in behavioral types: individual differences in behavioral responses remain stable with time and are consistent across various contexts. Such differences are referred to as ‘animal personalities’ (Roche, Careau, and Binning 2016), ‘temperaments’ (Réale et al. 2007), ‘coping styles’ (Koolhaas et al. 2007), or ‘endophenotypes’ (Gould and Gottesman 2006). When several different measures of behavioral traits correlate with each other in their individual expression then such between-individual correlations of behavioral traits are referred to as ‘behavioral syndromes’ or ‘personality dimensions’ (Sih et al. 2004; Réale et al. 2007; Dingemanse et al. 2012; Dingemanse and Dochtermann 2013; but also see Beekman and Jordan 2017). It has been suggested that different behavioral types can be maintained in natural populations via frequency-dependent selection because different behavioral responses have similar fitness payoffs but are better adapted to slightly different environmental conditions (Benus et al. 1991; Sih et al. 2004; Dingemanse and Réale 2005; Penke et al. 2007; Wolf et al. 2007; Wolf et al. 2008; Guilette et al. 2011). However, in addition to the genetic contribution to behavioral variability (Dochtermann, Schwab, and Sih 2015), individual differences may also arise from a state-behavior positive feedback loops (Sih et al. 2015), for example from conditions or events during brain maturation.

The idea that individuals tend to exhibit only a limited range of behavioral responses compared to the full behavioral repertoire of the species is interesting from an evolutionary perspective. On the other hand, models of decision making and resource use will remain incomplete if they do not take the constraints and consequences arising from individual differences into account.

Here we present a study on the foraging behavior of a population of wild nectar-feeding bats (*Glossophaga commissarisi* Gardner). Bats from the genus *Glossophaga* exhibit considerable individual variation in foraging behavior (Winter and Stich 2005; Thiele 2006). Compared to other members of the Glossophaginae subfamily (e.g. *Hylonycteris* or *Choeronycteris*) they are less specialized nectarivores with a broader diet (Tschapka 2004), which may result in a variety of foraging strategies adapted to different seasonal and local habitat conditions. Working with *Glossophaga soricina* in the laboratory we have performed pilot exploratory analyses searching for personality traits that satisfy the conditions of being different among individuals but stable with time. We have noticed that individuals on average maintain the same level of activity (daily number of visits to flowers) and tend to exploit a similar number of feeding locations (daily number of different flowers visited) over observation periods lasting for several months. Visit duration (time spent in hovering flight in front of or clinging to a flower) is another trait that showed individual consistency and it also tended to correlate negatively with activity, i.e. bats that make few visits make longer-lasting visits and vice versa. We hypothesized that the number of visits and visit duration might both be a part of a general *activity* personality dimension (Réale et al. 2007). The measure of different flower locations visited we interpreted as an indicator of how much an animal invests in information-gathering while foraging or, for the case of low numbers, its tendency to form behavioral routines. This can be considered as the exploration-exploitation balance in theoretical treatments of reinforcement learning (Daw et al. 2006). It must not be confused with the behavioral tendency to explore novel environments or objects. This latter concept is referred to as the *exploration-avoidance* continuum (Réale et al. 2007) i.e. the conflict between the motivation to explore novel contexts vs. the anxiety of an unknown that may be harmful. In our experiments bats were tested on a daily basis in a familiar environment, which eliminated the element of novelty. Furthermore, studies in rodents (Benus et al. 1991; Koolhaas et al. 2007; Coppens et al. 2010) and theoretical analyses of responsiveness (Wolf et al. 2008) indicate that routine formation and cue dependency are correlated with a tendency for aggression and that individual differences in aggressive behavior may reflect more general differences in how animals cope with environmental challenges. Consequently, the measure of number of different flowers visited can be taken as an indicator of an animal’s *proactivity-reactivity* coping style (Koolhaas et al. 2007; Coppens et al. 2010; or *shyness-boldness* personality dimension, Réale et al. 2007). However, in the absence of measures of other behaviors from this dimension, this interpretation is tentative.

We tested these hypotheses by investigating the foraging behavior of the species *G. commissarisi* in its natural habitat. The goals of this study were to: (*i*) confirm that previously identified behavioral traits in the foraging context exhibit consistency within individuals, (*ii*) investigate the potential link between personality traits and individual measures of foraging performance, and (*iii*) test whether the correlational patterns between these behavioral traits are consistent with previously hypothesized syndrome structures. We reanalyzed data from a field study in which free flying, individually tagged bats visited an array of computer operated flowers that provided nectar differing in sugar concentration (Nachev and Winter 2012). From the individual records of flower visitation, we determined the behavioral parameters *number of visits per bout*, *visit duration*, and *flowers visited during nightly foraging*. We then used these parameters to construct multi-response multivariate generalized linear mixed models (glmm), which allowed us to assess the within and between-individual correlations of the three parameters (Dingemanse and Dochtermann 2013). We used generalized linear mixed models to investigate whether foraging performance measures were correlated with these other behavioral measures. Finally, we compared hypothesized syndrome structures using the observed correlation pattern of the behavioral measures and deviance information criterion (DIC)-based model comparison (Dingemanse and Dochtermann 2013).

## METHODS

We analyzed the behavior of 51 adult *G. commissarisi* bats (21 females and 30 males) at La Selva Biological Station, Province Heredia, Costa Rica. The data were originally collected for a different study (Nachev and Winter 2012), and reanalyzed here because of the rarity of field data on a large number of individuals over a longer time period (over two months). Bats were caught by mist-netting, marked with 100 mg radiofrequency identification tags, measured, and released at the site of capture. As an indicator of a bat’s size, we measured forearm length with calipers. Bats had free access to a patch of artificial flowers - a rectangular array of 24 computer-controlled artificial flowers suspended horizontally under a steel frame canopy (Fig. 1 in Nachev et al 2017). The distance between flowers in the same row was about 40 cm and the distance between rows, about 60 cm. The flowers delivered rewards of 55-60 µL on every visit. Repeated visits to the same flower were always rewarded. Each artificial flower was connected via valves to two independent nectar pumps containing different concentration sugar solutions. This allowed us to program the experimental system such that half of the flowers provided a higher sugar concentration and the other half a lower sugar concentration. During a single night an individual flower’s concentration was fixed, but between nights, flower concentrations were systematically varied throughout the experiment. Thus bats had to relearn the quality of the flower locations each night. The data set consisted of series of two-alternative choice tests, with 12 flowers per option. The extent to which bats discriminate between two options of different food quality depends on both the difference between the two concentrations, and the average of the concentration pair (Nachev and Winter 2012; Nachev et al. 2013). Thus, for every night we calculated this relative intensity measure from the two sugar concentrations (Eq. 4 with *β* = 1.4 in Nachev et al. 2013). We then used this measure as a fixed predictor variable in further analyses.

**Fig. 1.**
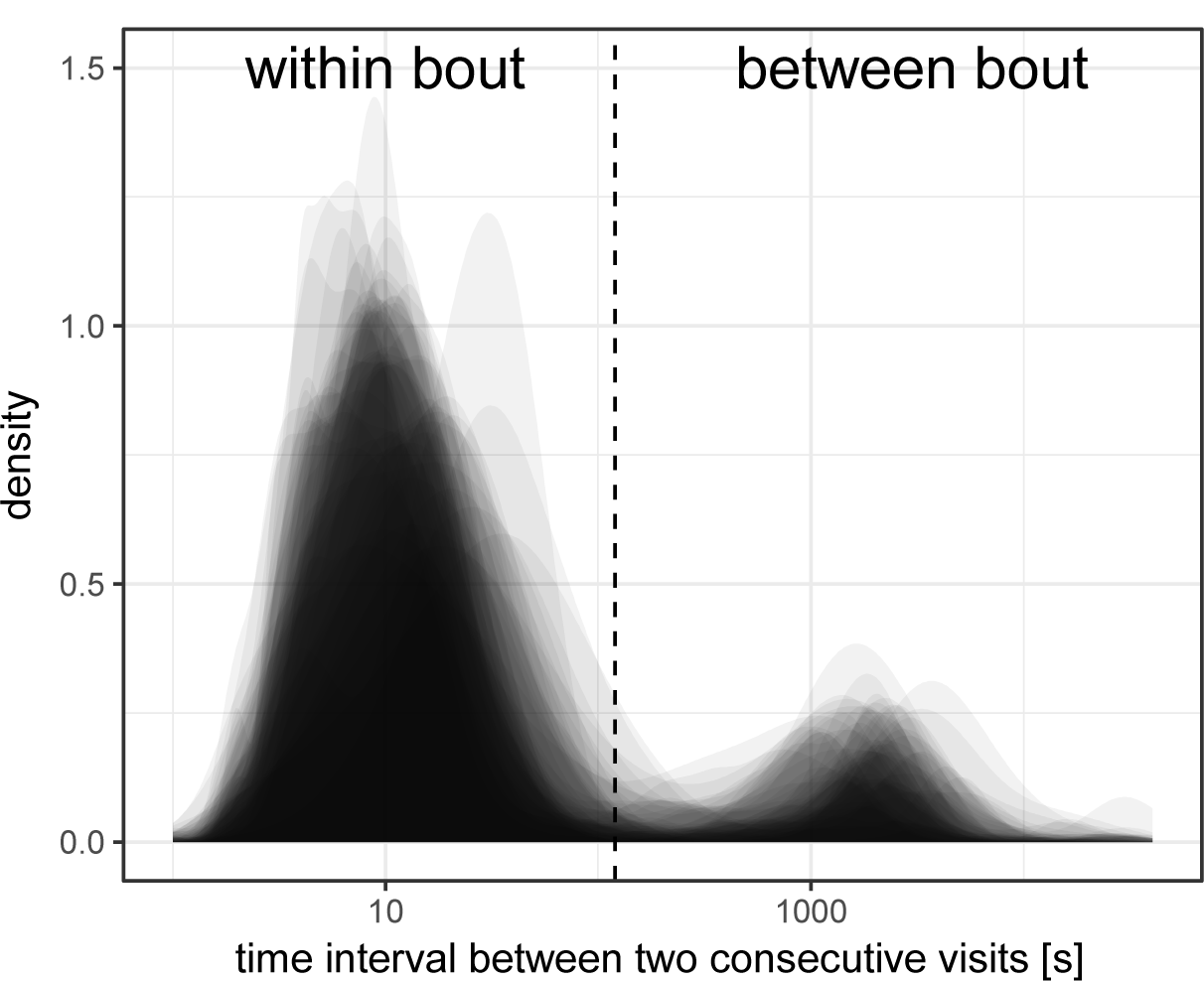
Bats’ foraging behavior was organized in clear bouts. The time intervals between two visits at a flower had a bimodal distribution when logarithmically transformed. Each transparent curve is based on the cumulative visits of an individual tagged bat. We selected the time interval of 120 seconds (vertical dashed line) as a criterion to distinguish visits within a bout from visits between bouts.

### Behavioral measures

The behavioral data collected during the experiments were the time stamped events (including starting time and duration) of individually identified bats visiting specific flowers that delivered rewards of known amount and sugar concentration. We present data from the 51 bats (out of a total number of 63 tagged bats) that made at least 24 flower visits per night on at least three nights. The number of repeated measures per focal individual was 22 ± 13 (mean ± SD; range: 3-44). Our sample of females consisted of 21 individuals, eight of which were pregnant. Due to the smaller sample size and potential confounding factor in females we mainly focus on the behavior of the 30 male individuals. Three behavioral measures were calculated for each bat for every experimental night. We analyzed the complete record of nightly events, in contrast to the analysis presented in Nachev and Winter (2012), where only events between 20:00h and 3:00h were analyzed. The behavioral measures we analyzed were:

**Visits per bout –** the total number of flower visits divided by the number of bouts made by an individual bat during a single night. We split the foraging behavior of bats into bouts using an interval of 120 seconds duration without flower visits as a bout break criterion (Figure 1). If a bat made fewer visits during a night than the number of flowers on the experimental array (*N* = 24), we excluded this day from the data set of this bat, in order to avoid a spurious positive correlation between number of visits per bout and number of flowers visited (see below).

**Visit duration –** the mean of all visit durations made by a bat during a single night. Longer durations of several seconds indicate a tendency to make clinging visits instead of the brief hovering visits, which normally lasted less than a second. However, there is no clear threshold duration value that separates the two behaviors.

**Flowers visited –** the total number of different flowers visited by an individual bat (ranging from 1 to 24) during a single night.

### Behavioral consistency

Based on prior experiments in the laboratory, we expected the individual differences between bats to remain stable over time. In order to test this, we performed repeatability analyses (Bell et al. 2009; Nakagawa and Schielzeth 2010) separately for males (*N* = 30) and females (*N* = 21). We used a multivariate random intercept and random slope model with *intensity* as a fixed effect and with visits per bout, visit duration, and flowers visited as independent variables (Hadfield 2009; Dingemanse and Dochtermann 2013; *MCMCglmm* package in R 3.4.3, R Development Core Team 2018). We used the Gaussian family for the parameters visits per bout and visit duration, and the Poisson family for flowers visited. For this, and each further analysis unless otherwise specified, we used 130,000 iterations with a thinning interval of 100 and a burn-in phase of 30,000, obtaining 1,000 samples for each estimate. Different priors (inverse Wishart, flat covariances, and parameter expanded; Hadfield 2009; Mutzel et al. 2013) did not result in qualitative changes in the output. From the output of these models we calculated the adjusted repeatabilities, that is the repeatabilities after controlling for confounding effects (Nakagawa and Schielzeth 2010).

From the same models we also obtained estimates for the between-individual and within-individual correlations of the three behavioral measures, along with their 95% credible intervals. We compared the observed between-individual correlation pattern with *a priori* hypotheses about behavioral syndrome structures (Figure 2; Dochtermann and Jenkins 2007; Dingemanse et al. 2010). Credible intervals that do not overlap with zero indicate statistical significance for paths between behavioral measures. We used credible intervals to distinguish between models 2 through 4 (Figure 2). Additionally, we tested if the three measures were independent from each other by running models in which the betindividual covariances were fixed at zero, but with the within-individual covariances left as free parameters. We compared the two models (model 1 and model 5, Figure 2) using the deviance information criterion (DIC; Spiegelhalter et al. 2002). A lower DIC score indicates higher explanatory power after penalizing for the number of free parameters. We considered models with ΔDIC>2 (i.e. with DIC scores differing by more than two from the model with the lowest DIC score) to be statistically unsupported.

**Fig. 2.**
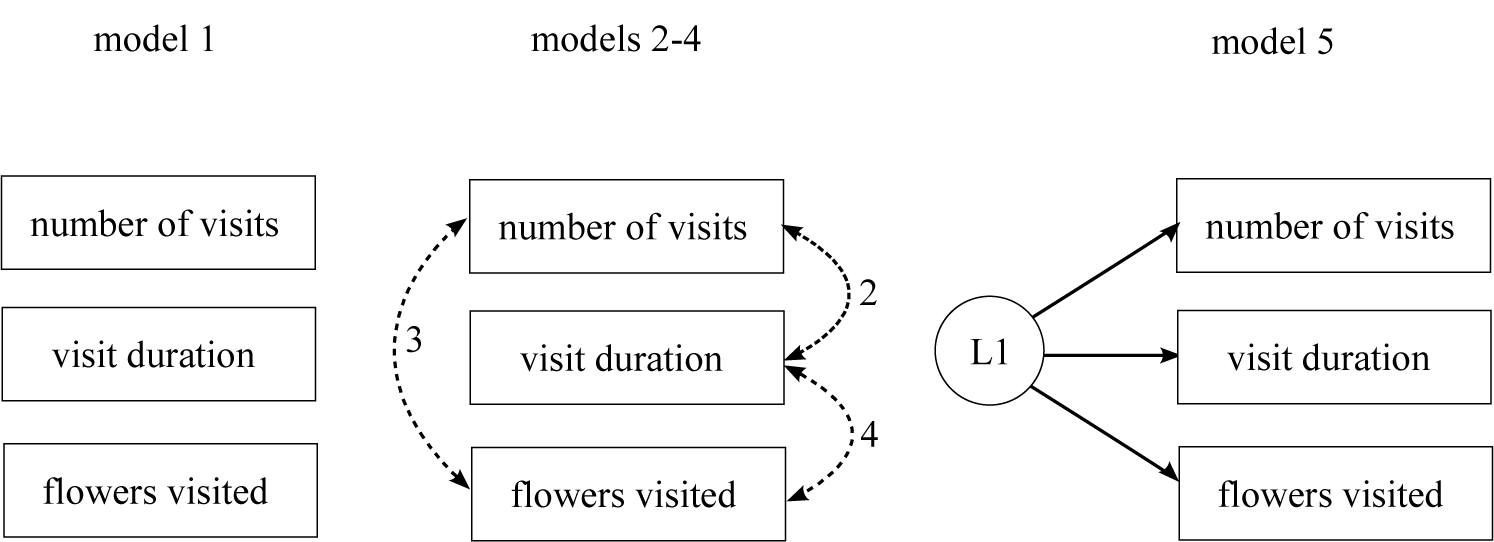
Models (1-5) of different syndrome structures. See *Methods* for model descriptions. Continuous unidirectional lines represent a causal relationship from a latent variable (L1) to behavioral measures. Dashed bidirectional arrows represent correlations between behavioral measures expressed in particular syndrome structures. In model 2 only path ‘2’ is active, in model 3 only path ‘3’ is active and in model 4 only path ‘4’ is active.

In order to test which of the three behavioral parameters visits per bout, visit duration, and flowers visited were influenced by intensity, sex, and body size, we also ran three univariate random intercept and random slope models with intensity, sex, and forearm length as fixed factors, and bat individual as random factor on the pooled data set for males and females (*MCMCglmm* package in R 3.4.3, R Development Core Team 2018). For these tests we corrected the *P* values using the Benjamini & Hochberg (a.k.a. false discovery rate) correction.

### Behavioral syndrome structure

*Model 1.* Behavioral independence. This was the null model with assumed lack of relationship between the three behavioral parameters.

*Model 2.* A link between visits per bout and visit duration (Figure 2, only path ‘a’ activated), both interpreted as indicators of the *activity* dimension (Réale et al. 2007). Flowers visited is independent from the other two measures.

*Model 3.* A link between visits per bout and flowers visited (Figure 2, only path ‘b’ activated). In this model visit duration is independent from the other variables and not an indicator of any particular personality dimension, whereas visits per bout and flowers visited both reflect the *activity* dimension.

*Model 4.* A link between flowers visited and visit duration and (Figure 2, only path ‘c’ activated), both interpreted as probable indicators of the *proactivity-reactivity* dimension (Koolhaas et al. 2007). The rationale behind this hypothesis is that an animal might make visits to a higher number of different flowers to sample their nectar concentration and abort the visits prematurely when the concentration is of the lower type. In this model visits per bout is the sole indicator of the *activity* dimension.

*Model 5.* Full domain-general syndrome with all measures indicators of the same dimension, (e.g. *activity*).

### Foraging performance

Foraging tasks are often modeled using temporal difference (TD) learning. Here, the expectations about finding a positive reward in an environment with resource locations of yet unknown quality are continually updated with ongoing experience. Choice performance then depends primarily on the learning rate (Glimcher 2011) and also the balance between exploration and exploitation, i.e. the sampling of locations with uncertain quality vs. the visitation of locations with a recently experienced positive outcome (Daw et al. 2006; Mathot et al. 2012). Consequently, we used two different measures of foraging performance, bouts to criterion and error rate (see below), both calculated from the seven nights on which the most extreme differences in sugar concentrations were presented to the bats (5% vs. 20%, 10% vs. 25%, and 15% vs. 30%). These pairs of sugar concentrations were chosen because they were associated with the highest, nearly perfect discrimination performance by bats (Nachev and Winter 2012). Under these conditions the effect of individual differences in perception of sugar concentrations is minimized, so that differences in learning rates and non-perceptual error rates (e.g. due to exploration) can be better estimated. For these measures the sample size was *N* = 23 bats, because we only analyzed males and not all bats were detected on more than two of these seven nights. Again, if a bat made fewer than 24 visits on a given night, this data point was not included.

**Visits to criterion –** the number of visits a bat made on a particular night, until the average proportion of visits to the higher sugar concentration flowers (discrimination performance) reached 0.8 or higher. In order to calculate this parameter, we used a change point algorithm (Gallistel et al. 2004; Nachev 2018) that separates a sequence of visits into chunks with significantly different choice preferences. We considered only chunks of at least ten visits. The visits to criterion measure was taken as the number of the first visit of the chunk in which the bat reached the criterion. Most males reached this criterion on every night on which they were detected, except for three bats, each on only a single night. Thus, the average number (± SD) of repeated measures per bat for this parameter in our data set was 5.7 nights ± 1.7. All else being equal, a high value for this measure indicates that a bat was faster in avoiding options with lower sugar concentrations and therefore gained a higher energy intake per visit. The probability of making a mistake and the overall number of visits needed to obtain a reliable estimate of the flower quality for each flower both scale with the total number of flowers visited. Therefore, we expected that bats visiting overall a smaller number of different flowers to reach criterion within a lower number of visits. On the other hand, we expected, all else being equal, for bats to reach the criterion just as fast, regardless of how they distributed their visits between bouts. In other words, we expected a bat making 50 visits in a single bout and a bat making ten bouts of five visits each (but visiting the exact same flowers in the exact same sequence) both to reach the criterion in the same number of visits. Therefore, we expected a lack of correlation between visits per bout and visits to criterion. Finally, we did not have a prior expectation for a relationship between visits to criterion and visit duration

**Error rate –** a measure of the relative frequency of errors (in this case, visits to low concentration flowers) due to factors of a non-perceptual nature, e.g. information-gathering or exploratory behavior. Error rate was calculated from the data after a bat had reached its nightly asymptotic choice behavior phase. The error rate is the proportion of visits to low concentration flowers after the last change point (Gallistel et al. 2004; Nachev 2018) in preference. We expected our stable experimental conditions (fixed sugar concentrations and volumes during a night) to favor bats with more routinized behavior. Once several advantageous flowers had been found a bat did not gain from further exploration. |Therefore, bats visiting only a few different flowers were expected to have lower error rates. Since we measured error rate from asymptotic behavior, we expected error rate to be unaffected by visits per bout. We had no prior expectation for the relationship with visit duration. As mentioned above, error rate was only determined from those experimental conditions where concentration differences were of high salience.

We ran another multivariate random intercept and random slope model as the one described in *Behavioral consistency*, but added visits to criterion and error rate as dependent variables (*MCMCglmm* package in R 3.4.3, R Development Core Team 2018). We used the Poisson family for visits to criterion and the multinomial family for the error rate. This allowed us to estimate the repeatabilities for the two measures of foraging performance, as well as the between and within-individual correlations for all behavioral measures. Because of the larger number of estimated parameters, we used 1,300,000 iterations with a thinning interval of 1000 and a burn-in phase of 30,000, obtaining 1,000 samples for each estimate.

### Data availability

The datasets generated during and/or analysed during the current study, including all statistical tests are available in the Zenodo repository: https://doi.org/10.5281/zenodo.2537944.

## RESULTS

Some 50–80 *G. commissarisi* bats foraged simultaneously during the course of the two-alternative choice experiment, making on average one visit per minute per individual. The average number (± SD; range) of flower visits by individual tagged bats per night was 2900 ± 1600 (400-9700). The three univariate glmm models revealed no significant differences between males and females in visit duration (estimate = -11.8, credible interval = - 41.0, 20.9, hereafter reported as -41.0 ≤ -11.8 ≤ 20.9, *P* = 0.55, all *P* values corrected for false discovery rate; Table 1). However, males made on average fewer visits per bout (-1.66 ≤ -0.95 ≤ -0.16, *P* = 0.024; Table 1) and visited fewer different flowers than females did (-1.67 ≤ - 1.55 ≤ -1.42, *P* = 0.005; Table 1). Forearm length did not significantly affect visits per bout (-0.53 ≤ -0.07 ≤ 0.32, *P* = 0.86), or flowers visited (-0.05 ≤ 0.00 ≤ 0.05, *P* = 0.94), but bats with longer forearms on average made visits with shorter durations (-35 ≤ -18 ≤ -1.3, *P* = 0.045). On average, visits per bout increased with the relative intensity of the difference in available concentrations (4.59 ≤ 9.74 ≤ 14.79, *P* = 0.005), but visit duration (-561 ≤ -288 ≤ -28.9, *P* = 0.072) or the number of flowers visited did not significantly depend on relative intensity of the sugar concentration difference (-1.20 ≤ 1.34 ≤ 2.46, *P* = 0.435).

**Table 1.** Phenotypic correlation (Pearson’s *r*) coefficients for behavioral parameters of the 51 bats.

|  | Visits per bout | Visit duration (ms) | Flowers visited |
| --- | --- | --- | --- |
| **Males ($N = 30$)** |  |  |  |
| Visits per bout | 7.73 (3.13)<sup>a</sup> |  |  |
| Visit duration (ms) | 0.05 (0.80)<sup>b</sup> | 659 (214) |  |
| Flowers visited | 0.33 (0.08) | **0.37 (0.04)** | 8.28 (3.09) |
| **Females ($N = 21$)** |  |  |  |
| Visits per bout | 8.86 (3.83) |  |  |
| Visit duration (ms) | **-0.64 (0.002)** | 662 (150) |  |
| Flowers visited | 0.18 (0.42) | 0.24 (0.29) | 13.10 (3.70) |

<sup>a</sup> Values on main diagonals give mean measures with standard deviations in parentheses.
These mean measures are calculated from the individual means.
<sup>b</sup> Correlations are based on individual means calculated over the whole data set, excluding nights in which bats made less than 24 visits, and bats that were detected on fewer than three nights. The corresponding two-tailed $p$ values are given in parentheses. Significant correlations are shown in bold.

The individual behavioral consistency of all three measures in the 51 tagged individuals was high, as indicated by the individual repeatability estimates for visits per bout (males: adj.*R* = 0.40, 95% credible interval = 0.26, 0.54, *N* = 30; hereafter reported as 0.26 ≤ 0.40 ≤ 0.54; females: 0.26 ≤ 0.39 ≤ 0.58, *N* = 21), visit duration (males: 0.59 ≤ 0.69 ≤ 0.81, *N* = 30; females: 0.37 ≤ 0.57 ≤ 0.70, *N* = 21), and flowers visited (males: 0.29 ≤ 0.46 ≤ 0.61, *N* = 30; females: 0.34 ≤ 0.49 ≤ 0.71, *N* = 21).

### Relationship between behavioral measures and measures of foraging performance

The two measures of foraging performance were both significantly repeatable within individuals (visits to criterion: 0.18 ≤ 0.30 ≤ 0.54, *N* = 23; error rate: 0.32 ≤ 0.49 ≤ 0.71, *N* = 23). The glmm revealed that bats that made more visits per bout on average took more visits to reach the criterion of 80% discrimination performance (Table 2). However, there were no other significant between-individual correlations between the two measures of foraging performance and any of the other three behavioral measures (Table 2). Within individuals, there was a significant negative correlation only between error rate and visit duration (Table 2).

**Table 2.** Between-individual and within-individual correlation coefficients for behavioral parameters and measures of foraging performance.

|  | Between individuals |  | Within individuals |  |
| --- | --- | --- | --- | --- |
| Males<br>(*N* = 23) |  |  |  |  |
|  | Visits to criterion | Error rate | Visits to criterion | Error rate |
| Error rate | -0.45 ≤ 0.05 ≤ 0.49 |  | -0.22 ≤ 0.13 ≤ 0.39 |  |
| Visits per<br>bout | **0.03 ≤ 0.58 ≤ 0.83** | -0.47 ≤ -0.09 ≤ 0.47 | -0.21 ≤ 0.11 ≤ 0.36 | -0.51 ≤ -0.11 ≤ 0.20 |
| Visit<br>duration | -0.53 ≤ 0.08 ≤ 0.45 | -0.61 ≤ -0.13 ≤ 0.35 | -0.31 ≤ -0.10 ≤ 0.19 | **-0.58 ≤ -0.35 ≤ -0.01** |
| Flowers<br>visited | -0.20 ≤ 0.21 ≤ 0.60 | -0.41 ≤ 0.05 ≤ 0.46 | -0.20 ≤ 0.32 ≤ 0.60 | -0.32 ≤ -0.02 ≤ 0.34 |

Note.- Correlations are based on 1000 posterior samples per individual. The values are given as lower 95% credible interval ≤ posterior mode ≤ upper 95% credible interval. Between-individual correlations are given to the left and within-individual correlations are given to the right. Values in bold do not overlap with 0.

### Analysis of behavioral syndrome structure

The between-individual correlations for the three behavioral parameters did not support any of the path connections between the different behavioral measures in males (models 2-4; Figure 2; Table 3), but in females there was support for model 2 (Figure 2; Table 2). The model in which the between-individual correlations were fixed to zero (model 1) and the model in which the correlations were left as free parameters (model 5) could not be distinguished from each other neither in males (model 1: DIC = 14627.03; model 5 DIC = 14626.55; ΔDIC = -0.48) nor in females (model 1: DIC = 11644.12; model 5: DIC = 11644.61; ΔDIC = 0.49).

**Table 3.** Between-individual and within-individual correlation coefficients for behavioral parameters of the 51 bats.

|  | Visits per bout | Visit duration | Flowers visited |
| --- | --- | --- | --- |
| **Males ($N = 30$)** |  |  |  |
| Visits per bout | | $-0.15 \leq -0.09 \leq 0.00$ | **$0.52 \leq 0.61 \leq 0.67$** |
| Visit duration | $-0.35 \leq 0.07 \leq 0.38$ | | $-0.04 \leq 0.09 \leq 0.17$ |
| Flowers visited | $-0.18 \leq 0.23 \leq 0.56$ | $-0.01 \leq 0.42 \leq 0.67$ | |
| **Females ($N = 21$)** |  |  |  |
| Visits per bout |  | **$0.03 \leq 0.13 \leq 0.21$** | **$0.49 \leq 0.59 \leq 0.66$** |
| Visit duration | **$-0.88 \leq -0.73 \leq -0.39$** |  | **$0.18 \leq 0.29 \leq 0.42$** |
| Flowers visited | $-0.42 \leq 0.11 \leq 0.48$ | $-0.24 \leq 0.07 \leq 0.59$ | |

Note.- Correlations are based on 1000 posterior samples per individual. The values are given as lower 95% credible interval ≤ posterior mode ≤ upper 95% credible interval. Between-individual correlations are given to the left and below the main diagonal and within-individual correlations are given to the right and above. Values in bold do not overlap with 0.

Even though there was no support for any of the between-individual correlations in males, there was a significant positive within-individual correlation between visits per bout and flowers visited (Table 3). In females all the within-individual correlations were significant and positive (Table 3).

## DISCUSSION

Consistent with our observations from the laboratory, the wild *G. commissarisi* in this study exhibited individual behavioral consistency in the number of visits per bout they made to a patch of artificial flowers, their mean visit duration, and the number of different flowers they visited. Except for visit duration, there was no evidence that the individual behavioral differences were related to differences in body size (from forearm length).

This individual behavioral consistency of the three parameters could not be explained by their interdependence within a full domain-general syndrome. We found no clear patterns of between-individual correlations, neither in males nor in females (Table 3). The mostly low and non-significant between-individual correlations suggest that the behavioral consistency most likely results from three independent dimensions. The only consistent finding in both male and female bats was that on nights on which bat individuals made a higher number of visits per bout, they were also more likely to visit a higher number of flowers. It was not the case, however, that bats that on average made more visits per bout also visited a higher number of different flowers on average, as indicated by the lack of significant between-individual correlations in either sex. The only significant between-individual correlation in females suggests that females either visit many flowers per bout but hovering very briefly in front of them, or they visit a smaller number of flowers per bout but make visits with longer durations. Surprisingly, the within-individual correlation between visits per bout and visit duration was significant, but in the opposite direction compared to the between-individual correlation. All in all, our findings demonstrate that phenotypic correlations between different behaviors (Table 1) are not necessarily indicators of behavioral syndromes (Table 3; Dingemanse et al. 2012; Dingemanse and Dochtermann 2013).

Without significant between-individual correlations between the three behavioral measures from our study and without knowing what other behaviors these correlate with, it is not possible to determine the mechanisms leading to the observed behavioral consistency. In the following, we provide some tentative interpretations and ideas for future studies. Concerning the pattern of resource exploitation, some bats consistently visited only a few of the available flowers, whereas others spread their activity over more than half of the flower array (range of mean number of flowers visited: 4-20; Table 1). We suggest that this difference may be a difference in the degree of behavior routinization (Koolhaas et al. 2007; see also Wolf et al. 2008). Our flowers always delivered rewards and the sugar concentrations of their nectars were stable within nights and only varied from night to night, thus favoring the development of stable choice patterns and penalizing unnecessary information-gathering. It remains to be shown that number of flowers visited is linked to known behaviors from the *proactivity-reactivity* (Koolhaas et al. 2007, e.g. aggressive interactions with conspecifics) or *shyness-boldness* continuum (responses to non-novel risk situations, Réale et al. 2007, e.g. delay to resume foraging after a perceived predator attack).

The distance bats travelled from their (night-time) roosts to the flower array is a potential uncontrolled confounding factor that could account for the repeatability of the number of visits per bout and potentially other behavioral measures. However, we first became aware of the repeatability of visits per bout in our laboratory studies, where all bats were kept in the same room with the flowers and considerations of travel distances were not applicable.

Our results provide some support to the hypothesis that different behavioral types may be better adapted to different environmental conditions (Guillette et al. 2011), more specifically, to different resource qualities and distributions. Though we did not assess fitness directly, differences in foraging efficiency can be positively correlated with fitness (Ritchie 1990; Lemon 1991; Jeanniard-du-Dot et al. 2017). The differences in both measures of foraging performance were significantly repeatable and the measure visits to criterion was significantly correlated with visits per bout, also a repeatable measure. Although this correlation was unexpected, we had based our prediction on the assumption that bats differing in visits per bout do not also differ in the spread of visits over the flower array *per bout*. However, if we define a new measure, flowers per bout, as the average number of different flowers a bat visits per bout during a single night, we can see that there is a significant correlation between flowers per bout and visits per bout (between-individual *r* = 0.20 < 0.62 < 0.80, within-individual *r* = 0.76 < 0.73 < 0.79). Thus, when bats make more visits within a bout, they tend to visit a higher number of different flowers, which seems to slow learning down and affect visits to criterion. Under conditions of higher reward uncertainty, we would expect bats that invest more in information-gathering to have better chances of detecting the locations of more profitable sources of nectar. On the other hand, especially at flowers with high nectar secretion rates, a more routinized behavior may confer fitness benefits through ‘defense by exploitation’ (Paton and Carpenter 1984; Ohashi and Thomson 2005). Foragers employing this strategy maintain high activity rates and exploit a limited number of replenishable food sources therefore keeping the mean resource standing crops low. This can reduce resource competition, because competitors using different strategies may perceive the shared food sources as unprofitable and leave to forage elsewhere. In our experiments the behavior of males seems consistent with defense by exploitation. Alternatively, the smaller numbers of visits per bout and flowers visited compared to females might also be an indication of agonistic interactions at the flowers.

Realistic models of bat foraging need to take into account the observed repeatable differences in frequency and distribution of flower visits. However, these differences also have important implications for the fitness of the plants that bats pollinate. For example, bats with different propensities to make revisits to the same plant or flower can exert very different selection pressures on plants, especially if the plants are self-incompatible. On the other hand, bats that make more visits per bout probably remove and deposit more pollen from flowers, since pollen can be ingested during grooming in the pauses between bouts. Finally, although hovering flight duration has been shown to be uncorrelated with pollen transfer (Muchhala and Thomson 2007), it may be that clinging visits result in higher pollen transfer than hovering visits. Thus, both the quality and quantity of pollination service provided by a pollinator may depend on its foraging strategy.

## Acknowledgments

We thank Arne Jungwirth for fieldwork assistance and Alexej Schatz for software programming. This work was supported by the National Geographic Society (8579-08); the Deutsche Forschungsgemeinschaft (Exc257, Exc277); and the Volkswagen Foundation (84915 to V.N.). This is a pre-print of an article published in Behavioral Ecology and Sociobiology. The final authenticated version is available online at: https://doi.org/10.1007/s00265-019-2637-4

## Conflict of Interest

The authors declare that they have no conflict of interest.

## Ethical Standards

Treatment of the experimental animals complies with the national laws on animal care and experimentation.

